# Knowledge Graph-based Thought: a knowledge graph enhanced LLMs framework for pan-cancer question answering

**DOI:** 10.1101/2024.04.17.589873

**Authors:** Yichun Feng, Lu Zhou, Chao Ma, Yikai Zheng, Ruikun He, Yixue Li

## Abstract

**Background:** In recent years, Large Language Models (LLMs) have shown promise in various domains, notably in biomedical sciences. However, their real-world application is often limited by issues like erroneous outputs and hallucinatory responses.

**Results:** We developed the Knowledge Graph-based Thought (KGT) framework, an innovative solution that integrates LLMs with Knowledge Graphs (KGs) to improve their initial responses by utilizing verifiable information from KGs, thus significantly reducing factual errors in reasoning. The KGT framework demonstrates strong adaptability and performs well across various open-source LLMs. Notably, KGT can facilitate the discovery of new uses for existing drugs through potential drug-cancer associations, and can assist in predicting resistance by analyzing relevant biomarkers and genetic mechanisms. To evaluate the Knowledge Graph Question Answering (KGQA) task within biomedicine, we utilize a pan-cancer knowledge graph to develop a pan-cancer question answering benchmark, named the Pan-cancer Question Answering (PcQA).

**Conclusions:** The KGT framework substantially improves the accuracy and utility of LLMs in the biomedical field. This study serves as a proof-of-concept, demonstrating its exceptional performance in biomedical question answering.

**Key Points:**

- We introduce a framework combining LLMs with KGs to improve factual accuracy in LLM reasoning.
- Our system is a flexible architecture that seamlessly integrates various LLMs.
- Utilizing a pan-cancer knowledge graph, we have proposed the first KGQA benchmark in the field of biomedicine.
- Case studies reveal our method enhanced LLMs in addressing biomedical challenges such as drug repositioning, resistance research, individualized treatment, and biomarker analysis.
- The method performs favorably in comparison to existing methods.

## Introduction

With the increasing prominence of Large Language Models (LLMs) in the field of artificial intelligence, the advent of influential models such as ChatGPT [1] and Llama [2] consequently catalyze the development of a wide array of applications in biomedicine and healthcare. However, LLMs still face the challenge of factual hallucination, where they generate incorrect statements due to limited inherent knowledge [3]. Factual hallucination presents a significant challenge for the practical use of LLMs, especially in real-world scenarios where factual accuracy is crucial. Consequently, there is a growing focus on addressing factual hallucinations in LLMs within the field of Natural Language Processing (NLP) [4, 5].

LLMs often struggle to capture and access factual knowledge, primarily due to three aspects: the inability to comprehend questions due to the lack of contextual information, the insufficient knowledge to generate accurate answers, and the incapacity to recall specific facts [6]. Consequently, researchers consider the fine-tuning technique as a solution to address these issues. For example, MedAlpaca [7] builds upon medical data to fine-tune Stanford Alpaca for applications related to medical question-answering and dialogue. ChatDoctor [8] is designed to simulate a conversation between a doctor and a patient by fine-tuning LLaMA with medical literature. Additionally, Med-PaLM [9] shows promising performance on the MedQA exam based on clinical corpora and human feedback. Meanwhile, aiming at the Chinese medical domain, LLMs such as BenTsao [10], DoctorGLM [11], and HuatuoGPT [12], are developed on the Chinese medical dialogue data. More recently, Zhongjing [13] and ChiMed-GPT [14] adopted full pipeline training from pre-training, SFT, to Reinforcement Learning with Human Feedback (RLHF) [15]. While fine-tuning can reduce hallucinations in large language models (LLMs), it brings about considerable training expenses. Additionally, it poses a critical challenge known as catastrophic forgetting. This issue manifests when a model forgets its previously learned information as a consequence of parameter modifications during the acquisition of new tasks. This forgetfulness results in a deterioration of performance on prior tasks, consequently constraining the model’s practical applicability [16, 17].

In addition to fine-tuning, researchers also enhance the output of LLMs through the field of prompt engineering. Prompt engineering focuses on the creation and optimization of prompts to improve the effectiveness of LLMs across various applications and research domains [18]. It can enhance the capabilities of LLMs in a wide range of complex tasks, including question answering, sentiment classification, and common-sense reasoning. Chain-of-thought (CoT) prompts [19] enable complex reasoning capabilities by incorporating intermediate reasoning steps. The Automatic Prompt Engineer (APE) proposes an automatic prompt generation method aimed at enhancing the performance of LLMs [20]. Prompt engineering offers a straightforward approach to harnessing the potential of LLMs without fine-tuning.

On the other hand, Knowledge Graphs (KGs) are repositories of vast quantities of high-quality structured data, offering the potential to effectively mitigate the issue of factual hallucinations when integrated with LLMs. Hence, employing KGs for question-answering can enhance the precision of the responses and furnish a dependable foundation for the factual verification of information produced by LLMs. Knowledge Graph Question Answering (KGQA) has long been a hot research topic. Before the advent of LLMs, certain studies [21, 22, 23] typically begin by retrieving a subgraph related to the question to reduce the search space, then perform multi-hop reasoning on this basis. This retrieval-plus-reasoning paradigm has shown its advantages over direct reasoning across the entire KG [24, 25]. Additionally, Researchers tackle KGQA by parsing the question into a structured query language (e.g., SPARQL) and using a query engine to obtain accurate answers [26, 27]. UniKGQA [28] introduces a unified fine-tuning framework for retrieval and reasoning, more closely linking these two stages. However, traditional KGQA methods usually perform poorly in accurate semantic understanding and high-quality text generation due to the lack of LLMs for retrieval and reasoning. Hence, recent research is increasingly utilizing external KGs to enhance LLMs in addressing KGQA challenges. For instance, StructGPT [29] navigates through knowledge graphs by identifying pathways from an initial seed entity to the target answer entity, while Think-on-Graph (ToG) [30] introduces iterative exploration of the knowledge graph, which can become inefficient with very large KGs. Additionally, RoG [31] necessitates fine-tuning to accurately generate and plan the relation paths. KG-GPT [32] opts for retrieving an entire subgraph from the knowledge graph and then deduces the answer through inference. Although these methods have achieved gratifying results in general areas, as shown in Figure 1(B), when the intermediate entity in the multi-hop question is unknown, it is impossible to retrieve the appropriate knowledge from the KG.

**Figure 1.**
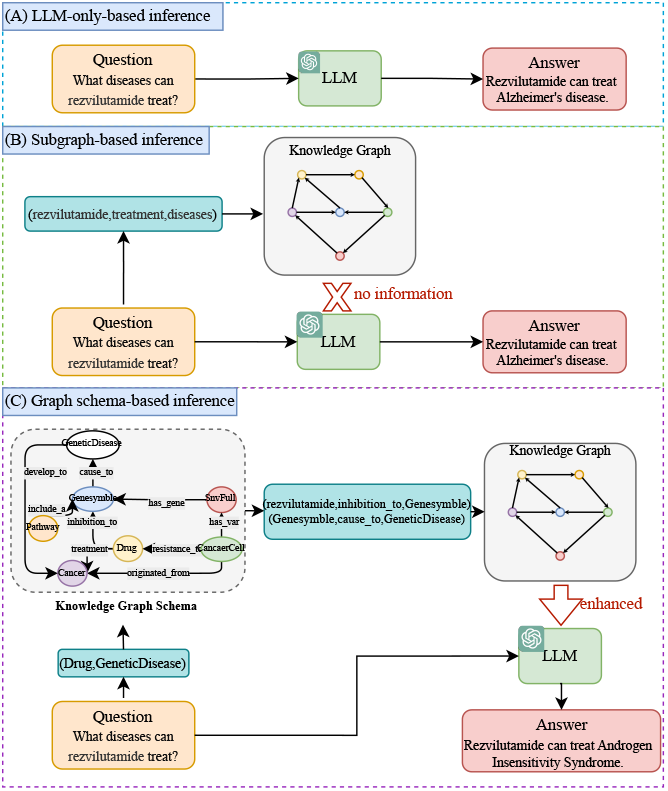
Illustrative examples contrasting our work with previous efforts. **(A) LLM-only-based inference**, answering questions solely through the inherent knowledge of LLMs. **(B) Subgraph-based inference**, enhancing LLMs by retrieving the knowledge from KGs based on the question. If intermediate entities are not provided in the multi-hop question, no appropriate knowledge can be retrieved. **(C) Graph schema-based inference**, enhancing retrieval capabilities by reasoning intermediary entity types on the schema of the KG, using the knowledge of the KG to enhance LLMs’ responses.

In this paper, we introduce an innovative framework called Knowledge Graph-based Thought (KGT), which integrates LLMs with KGs through employing LLMs for reasoning on the schema of KGs to mitigate factual hallucinations of LLMs, as shown in Fig. 1(C). Unlike traditional methods, KGT does not directly retrieve factual information based on the question. Instead, it uses LLMs to infer entity information on the schema of the knowledge graph, generating an optimal subgraph based on key information directly extracted from the question and inferred information from the schema. Subsequently, the optimal subgraph is used to infer the answer to the question through LLMs. KGT requires no fine-tuning, offers seamless integration with multiple LLMs, and is plug-and-play, facilitating easy deployment. It demonstrates generalizability, making it adaptable for use with diverse knowledge graphs. This framework is tailored for wide-ranging applications in numerous biomedical challenges, such as: (1) enhancing clinical decision-making for physicians and medical organizations; (2) delivering medical advice to patients and healthcare providers; (3) uncovering crucial biomarkers for early disease detection and tailored therapy; and (4) exploring novel therapeutic applications for existing medications through insights into their mechanisms, side effects, and the biological processes of associated diseases. Furthermore, we utilize the SmartQuerier Oncology Knowledge Graph (SOKG), a pan-cancer knowledge graph developed by SmartQuerier, to create a benchmark for the Knowledge Graph Question Answering task within biomedicine, named the Pan-cancer Question Answering (PcQA). We release this benchmark and its accompanying knowledge graph, which is a subgraph of the SOKG, in [33]. This benchmark is currently the sole question answering dataset available in the domain of biomedical knowledge graphs.

## Materials and Methods

### Knowledge graph introduction

In this work, we tackle the problem of logical reasoning over the KG 𝒦 : *E* × *R* that store entities (*E*) and relations (*R*). Without loss of generality, KG can be organized as a set of triplets {(*e*_1_, *r, e*_2_)} ⊆ 𝒦, where each relation *r* ∈ *R* exists between the pair of entities (*e*1, *e*2) ∈ *E* × *E*. We define a relational path {(*t*1, *r, t*2)} as a sequence of entity types (*T*) and the relation between them, where (*t*1, *t*2) ∈ *T* × *T*. In contrast, a relational chain {(*e*1, *r, e*2)} refers to a specific set of relational triplets between entities. To further enrich the KG, attribute information is included through pairs (*e, attr*), where *attr* represents an attribute associated with an entity *e*, thereby enhancing the KG’s semantic richness and precision by incorporating detailed characteristics of each entity.

Within the specialized realm of pan-cancer research, we use a subgraph of the SOKG that provides detailed oncological information. As depicted in Table 1, SOKG includes a collection of over 3 million entities, which is substantially larger than the entity count in the compared knowledge graphs, SynLethKG [34] and SDKG [35], with 540,012 and 165,062 entities, respectively. Furthermore, SOKG’s nearly 6 million unique concept relations exceed those of SynLethKG and SDKG, which have 2,231,921 and 727,318 relations, respectively. Additionally, SOKG includes 98 distinct attribute types, enriching data comprehension and improving the efficiency and precision of queries, a capability not matched by SynLethKG or SDKG, which do not include comparable attributes. For this research, we utilize only a subgraph of the SOKG, which is available as open data [33], while the full knowledge graph remains proprietary.

**Table 1.** Comparison of SOKG with SynLethKG and SDKG.

|  | Entity Types | Relational Types | Nodes | Edges | Attributes |
| --- | --- | --- | --- | --- | --- |
| SynLethKG | 11 | 24 | 54,012 | 2,231,921 | 0 |
| SDKG | 7 | 12 | 165,062 | 727,318 | 0 |
| SOKG | 24 | 21 | 3,640,259 | 10,656,273 | 98 |

### Tasks description

In order to tackle a diverse array of challenges in the field of biomedicine, we have designed four categories of problems: one-hop problems, multi-hop problems, intersection problems, and attribute problems, as illustrated in Table 2. Based on these four types of tasks, we leverage the SOKG to establish a benchmark for the Knowledge Graph Question Answering task within biomedicine, named the Pan-cancer Question Answering (PcQA). Unlike KGQA tasks in general domains, such as MetaQA[36] and FACTKG[37], which typically provide the entity types of intermediate entities, KGQA problems in the biomedical domain often do not have any information about intermediate entities. Instead, the information about intermediate entities must be inferred from the question itself rather than being directly provided as shown in Supplementary Material Table.S1. Additionally, our PcQA dataset includes attributes such as whether a drug is targeted therapy or if a mutated gene is oncogenic. This makes our tasks slightly more challenging and better suited to the actual needs of biomedical KGQA.

**Table 2.** Four different reasoning types of task. Each reasoning type may include overlapping questions, so the sum across the four different reasoning types of the task may exceed the total number of questions.

| Reasoning Type | Claim Example | Graph | Question Number |
| --- | --- | --- | --- |
| One-hop | What types of cancer can be treated with diethylstilbestrol? | $H_1 \xrightarrow{R_1} T_1$ | 243 |
| Multi-hop | What genetic mutations are present in adenoid cystic carcinoma? | $H_1 \xrightarrow{R_1} M \xrightarrow{R_2} T_1$ | 124 |
| Intersection | Which drugs are ALK in basaloid large cell carcinoma of the lung sensitivity to? | $H_1 \xrightarrow{R_1} T_1$<br>$H_2 \xrightarrow{R_2} T_2$<br>$T_1 \cap T_2 \rightarrow T_?$ | 37 |
| Attribute | What is the maximum age for recruitment of clinical trials for patients with meningioma? | $H_1 \xrightarrow{R_1} P_{T_1}$ | 59 |

#### One-hop problems

One-hop problems involve single-relation chain reasoning, where the objective is to deduce the tail entity *T*? given a head entity *H*_1_ and a relation *R*1, or to infer the relation *R*? when a head entity *H*1 and a tail entity *T*_1_ are known, as depicted in Equ.1 and Equ.2.

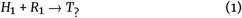

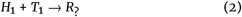

#### Multi-hop problems

Multi-hop problems involve multiple-relation chain reasoning, that can be broadly categorized into two types. The first category involves deducing potential relationships between entities by navigating through indirect relations. By examining the indirect relations (*R*1,*R*2) between a head entity *H*1 and a tail entity *T*1, it is possible to infer an unknown or potential relation *R*? linking them directly. This inference process is encapsulated in the following equation:

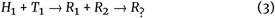

The second category extends the reasoning to include the discovery of entities themselves, by following a path from a head entity through intermediate relations to a final tail entity. Starting with a head entity *H*_1_, coupled with an indirect relation *R*_1_, an intermediary entity *M* can be inferred. This intermediary entity *M* is then applied with an indirect relation *R*_2_ to deduce the final tail entity *T*?. This inference process is summarized in the following equation:

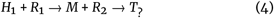

#### Intersection problems

Intersection problems refer to taking the intersection of multiple relational chains. Two head entities (*H*_1_,*H*_2_) lead to the deduction of two types of tail entities (*T*1,*T*2) based on different relations (*R*1,*R*2). The final tail entity *T*? is determined by intersecting these two types of tail entities (*T*_1_,*T*_2_). This inference process is summarized as following:

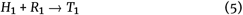

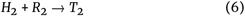

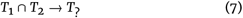

#### Attribute problems

Attribute problems refer to the attribute information of entity, where the task involves retrieving the attributes of a known head entity *H*_1_ or determining whether the tail entity *T*_1_, identified through a known head entity *H*_1_ and relation *R*_1_, satisfies the attributes specified in the query, as illustrated in Equ.8 and Equ.9.

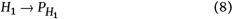

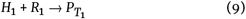

### Datasets

In the continuously evolving field of biomedical research, the integration of LLMs with KGs offers a more efficient and effective method for knowledge discovery and utilization, particularly in advancing cancer research. Nonetheless, we note a scarcity of appropriate datasets for evaluating these sophisticated methodologies within this field. To address this, we leverage the SOKG to establish a benchmark for the Knowledge Graph Question Answering task within biomedicine, named the Pan-cancer Question Answering (PcQA). Our questions were carefully crafted by experts based on the content of the knowledge graph. GPT-4 [38] was then employed to generate Cypher queries, which were used to retrieve answers from the knowledge graph. The generated Cypher queries and corresponding answers underwent an initial review by a biomedical PhD candidate, who manually verified and corrected the dataset against the knowledge graph. Finally, the entire dataset was thoroughly reviewed by two biomedical experts to ensure its accuracy and reliability. This multi-step process was meticulously designed to uphold the highest standards of quality throughout the dataset creation. This dataset, along with the accompanying knowledge graph, is completely open-source[33]. The PcQA includes 405 data entries, covering a wide range of applications in the field of pan-cancer research, including genetic predisposition to cancer, medication treatment planning, drug repositioning, identification of potential drug targets, studies on drug resistance, and predictions of cancer progression and metastasis. By deeply exploring cancer-related reasoning and information retrieval challenges, this dataset can inspire researchers and clinicians to gain a deeper understanding of cancer and explore more effective treatment methods.

### KGT framework

The overall framework of KGT is laid out in Fig. 2. When users input their question in natural language, the first step is to analyze the question, extracting the main information with the goal of breaking down the question into smaller, more manageable units. This main information is then passed to a LLM, which applies graph reasoning on the schema graph of the knowledge graph, yielding the optimal relational path. Subsequently, a retrieval statement is generated, and a subgraph is constructed within the KG through search. The relational chains and attributes in the subgraph are then fed back into the LLM to finalize the reasoning and generate an output in natural language.

**Figure 2.**
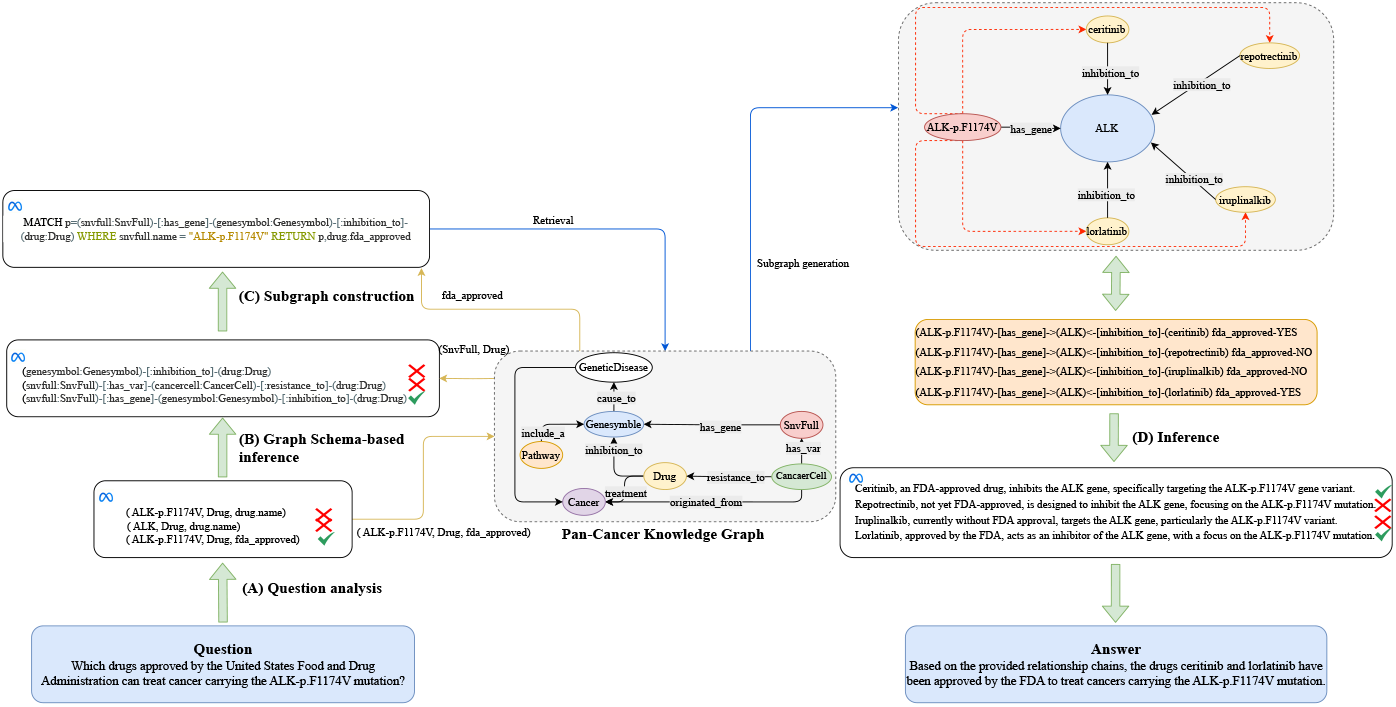
Framework of KGT. (A) Question analysis. Decompose the question and extract its key information. (B) Graph Schema-based inference. Input the types of the head and tail entities into the graph schema of the knowledge graph, complete the graph reasoning, and obtain the optimal relational path. (C) Subgraph construction. Generate a query statement and retrieve the subgraph. (D) Inference. Complete the final reasoning and output the results in natural language. Note: The symbol “*×*” represents content that has been filtered out by the LLM, while “✓” denotes the optimal content selected by the LLM.

#### Question analysis

##### Key information extraction

The user inputs a question text (*Q*) in natural language, which is initially deconstructed and parsed. A LLM is applied to analyze the question, resulting in the identification of the head entity name (*Hn*), the tail entity type (*Tt*), and the attributes of tail entity (*Ta*). The prompt for the LLM to extract key information from the question is presented in Supplementary Material Fig.S1.

##### Retrieving key information from KG

Based on *Hn*, a fixed Cypher format is set to query the head entity type (*Ht*), facilitating subsequent reasoning.

#### Graph Schema-based inference

#### Construction of a graph based on KG schema

Based on the entity types (*Et*) and the relations (*R*) between them in the SOKG, an undirected graph 𝒢 is established where *Et* serve as nodes 𝒩 and *R* act as edges 𝒫.

##### Candidate Path Search

Breadth-First Search (BFS) is employed to identify the shortest paths connecting *Ht* and *Tt* from the constructed graph 𝒢. Initiate the search at *Ht*, creating a queue to hold nodes encountered along the way. Simultaneously, form a set to track nodes that have been visited to avoid revisiting them. Insert *Ht* into the queue. Continue processing as long as the queue remains non-empty, removing a node from the queue at each step. For each of its unvisited neighbors, enqueue the neighbor, mark it as visited, and log the pathway from *Ht* to this neighbor. Upon arrival at *Tt*, use the accumulated path data to compile the set of shortest paths (*SPs*) from *Ht* to *Tt*, with each individual path within the set referred to as an *SP*. The nodes in each *SP* represent entity types, while the edges denote the relationships between these entity types.

##### Optimal path selection

By utilizing embedding technology, textual information is mapped into a low-dimensional space, resulting in N-dimensional real-value vectors. The similarity between each *SP* and the *Q* is calculated based on their respective real-value vectors, with the *SP* exhibiting the highest similarity being selected as the optimal path (*OP*).

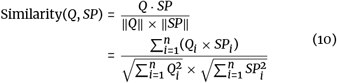

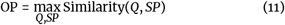

#### Subgraph construction

##### Query statement generation

Input *Ht, Hn, Tt, Ta*, and *OP* into an LLM to generate a query statement, such as Cypher. Text2Cypher Prompt is presented in Supplementary Material Fig.S2.

##### Subgraph generation

Enter the query statement in the KG to obtain a reasonable subgraph.

#### Inference

##### Subgraph inference

Based on the relational chains and attribute data in the subgraph, determine the relevance to the question text. Prune any erroneous information, retaining only the correct relational chains.

##### Natural language output

The LLM divides the subgraph into multiple relational chains, each of which outputs a sentence in natural language, and then the LLM generates natural language output. LLMs Inference and Output Prompt is presented in Supplementary Material Fig.S3.

## Results

### Evaluation criteria

We use evaluators based on GPT-4 [38], BERTScore [39], and ROUGE [40] to assess the accuracy of the generated answers. As a scoring bot, GPT-4 evaluates and assigns scores based on the similarity in meaning between two sentences. GPT-4-based Evaluation Prompt is presented in Supplementary Material Fig.S4. BERTScore evaluates semantic similarity using context-sensitive embeddings, offering a comprehensive evaluation of language model outputs. ROUGE, on the other hand, evaluates the longest common subsequence (LCS) between the generated text and the reference text, focusing on sequence-based similarity to assess the fluency and the preservation of semantic content.

### Baselines

To assess the advantages of our framework, we compare it with several approaches that can be directly applied for KGQA tasks without fine-tuning. We introduce a straightforward baseline approach, named Base, which is similar to KG-GPT [32], currently the leading method in the KGQA field, excluding the sentence segmentation step of KG-GPT. Initially, this involves leveraging a LLM to retrieve relevant information from the KG by generating a query statement. Then, another LLM is used to answer the question with the retrieved information. To enhance the baseline, we incorporate Chain-of-Thought (CoT) prompting [19] and In-Context Learning (ICL) techniques [41], collectively referred to as CoT&ICL. The prompts for these methods are illustrated in Supplementary Material Table.S5. Additionally, we implement KG-GPT [32] to enhance the retrieval and reasoning capabilities of the LLMs. For a fair comparison, all methods are based on Code-Llama-13B [42].

To further underscore the efficacy of our framework, we conduct a comparative analysis of KGT, which is built upon Code-Llama-13B, against two highly capable large language models that are prominent in the general and biomedical domains: ChatGPT-3.5 [1] and Taiyi [43]. ChatGPT-3.5, a leader in tasks across the general domain, has exhibited competitive performance in a wide range of applications. To compensate for its limited biomedical knowledge, we employed two methodologies previously described, Base and CoT&ICL, as advanced baselines to augment ChatGPT-3.5’s capabilities. Taiyi, a cutting-edge LLM in biomedicine, pre-trained on two trillion tokens, leverages its extensive biomedical knowledge base for direct question answering, bypassing the need for knowledge graph retrieval.

Due to the scarcity of KGQA datasets within the biomedical domain, all experiments are conducted on our newly proposed benchmark, named PcQA.

### Comparative analysis across different KGQA methods

We evaluated the capabilities of various methods based on Code-Llama-13B, with the experimental results presented in Table 3. The experimental results indicate that the Code-Llama-13B model, enhanced with KGT, consistently surpasses competing methods across all metrics assessed. Notably, KG-GPT improves the F1 score by 15.7% over previous methods CoT&ICL, while our method KGT increases the F1 score by 33% over KG-GPT. Because KG-GPT overlooks the impact of entity types and attributes on answers within the biomedical domain. This achievement positions our approach as a pioneering benchmark in biomedical KGQA, eclipsing previously established best practices.

**Table 3.** Comparison of results between KGT and other commonly used methods based on the Code-Llama-13B. Display the best results in bold for each indicator.

| Method | GPT-4 Eval (%) | BERTScore (%) | ROUGE (%) |  |  |
| --- | --- | --- | --- | --- | --- |
|  |  |  | Recall | Precision | F1-score |
| Base | 46.6 | 85.3 | 25.3 | 28.5 | 24.5 |
| CoT&ICL | 57.9 | 88.8 | 38.9 | 39.4 | 37.6 |
| KG-GPT | 68.2 | 93.5 | 55.2 | 55.8 | 53.3 |
| **KGT(ours)** | **92.4** | **97.7** | **87.4** | **87.7** | **86.8** |

### Comparative analysis across diverse LLMs

We presents a comparative study of KGT applied to Code-Llama-13B against two highly capable LLMs in the general and biomedical domains, with experimental results displayed in Table 4. Code-Llama-13B, enhanced by KGT, significantly outperforms its peers, achieving the highest marks in every assessment metric: a GPT-4 Eval score of 92.4, a BERTScore of 97.7, and a ROUGE F1-score of 86.8. Remarkably, our approach’s F1 score surpasses that of ChatGPT-3.5 with the Base method by 52.7%, the CoT&ICL method by 36.3%, and Taiyi’s base model by 67.3%. These results highlight KGT’s substantial contribution to improving the performance of large language models for the pan-cancer KGQA task. Even when integrated with open-source general models, KGT exhibits remarkable performance, outstripping both the recognized state-of-the-art closed-source large language models and those specifically tailored for the biomedical domain. This showcases KGT’s adeptness at parsing and leveraging knowledge graph data, setting a new standard for future research and applications in the field.

**Table 4.** Comparison of KGT based on Code-Llama-13B with results from other commonly used models. Display the best results in bold for each indicator.

| Model | Method | GPT-4 Eval (%) | BERTScore (%) | ROUGE (%) |  |  |
| --- | --- | --- | --- | --- | --- | --- |
|  |  |  |  | Recall | Precision | F1-score |
| ChatGPT-3.5 | Base | 65.4 | 91.0 | 42.7 | 32.3 | 34.1 |
|  | CoT&ICL | 70.3 | 93.3 | 57.0 | 50.6 | 50.5 |
| taiyi | \ | 40.6 | 85.3 | 15.4 | 39.6 | 19.5 |
| **Code-Llama-13B** | **KGT(ours)** | **92.4** | **97.7** | **87.4** | **87.7** | **86.8** |

### Assessing KGT’s effectiveness on diverse LLM platforms

To underscore the adaptability and effectiveness of our KGT framework when applied to a range of large language models, we conduct experiments on several LLMs: Zephyr [44], Llama-2 [2], and Code-Llama [42]. The outcomes, illustrated in Fig. 3, reveal that while the CoT&ICL techniques significantly boost performance in terms of F1-score, our KGT methodology delivers even more substantial enhancements across all evaluated models. This demonstrates not only the effectiveness of CoT&ICL as a performance-enhancing strategy but also highlights the superior advancements and impact of KGT, establishing its dominance and efficiency in knowledge graph question-answering tasks.

**Figure 3.**
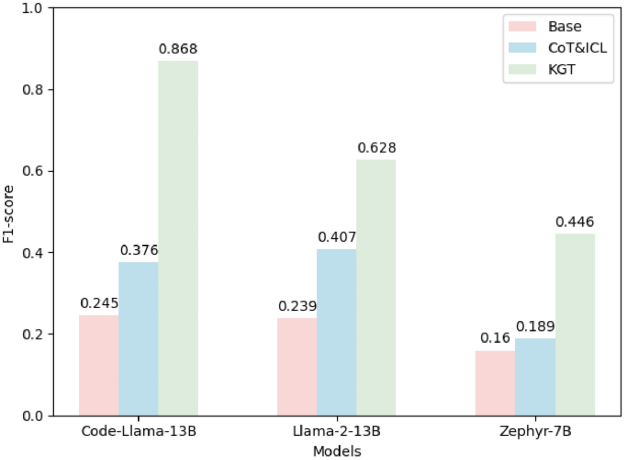
Performance of various models using different strategies.

### Ablation study for dissecting the components of KGT

In our effort to illuminate the individual contributions of the components that constitute our KGT framework and their collective impact on enhancing the performance of LLMs, we define four foundational modules: (1) question analysis for the extraction of pivotal information, (2) graph schema-based inference to identify the optimal relational chains in the knowledge graph, (3) the generation of query statements to facilitate subgraph construction, and (4) the inference process coupled with the articulation of results in natural language. This ablation study, grounded on the Code-Llama-13B model, is meticulously designed to evaluate the efficacy of these components. Since graph schema-based inference requires the process of question analysis, the question analysis module cannot be removed in isolation; simultaneously, subgraph construction is indispensable for knowledge graph retrieval. If the subgraph construction module is independently omitted, the outputs of the initial two modules will not impact the final results, making the isolated exclusion of this component illogical. Therefore, we introduce three specific ablated configurations for examination: (1) excluding graph schema-based inference (*w/o* GSBI), (2) omitting both question analysis and graph schema-based inference (*w/o* QA&GSBI), and (3) removing question analysis, graph schema-based inference, and subgraph construction (*w/o* QA&GSBI&SC), effectively bypassing the structured query of the SOKG and relying solely on the LLM’s inherent knowledge for question answering.

The results of the ablation study, as shown in Table 5, demonstrate that when we remove the GSBI, we observe a 20% decrease in the F1 score. Removing both GSBI and QA results in an additional 8.6% decrease in the F1 score compared to removing GSBI alone. Furthermore, removing GSBI, QA, and SC together leads to a 46% decrease in the F1 score compared to removing just GSBI and QA. The experiments reveal that SC is crucial; its absence forces the LLM to rely solely on its inherent knowledge, significantly reducing effectiveness. GSBI is also key, as it aids in navigating complex multi-hop questions by providing necessary intermediate entity information for subgraph construction. QA is equally essential, ensuring accurate identification of entities and properties for correct subgraph construction. All these variants underperform compared to the complete KGT, indicating that each of the three modules is vital for the final performance. Furthermore, such observations confirm that our KGT can indeed leverage knowledge to enhance the final performance of LLMs.

**Table 5.**
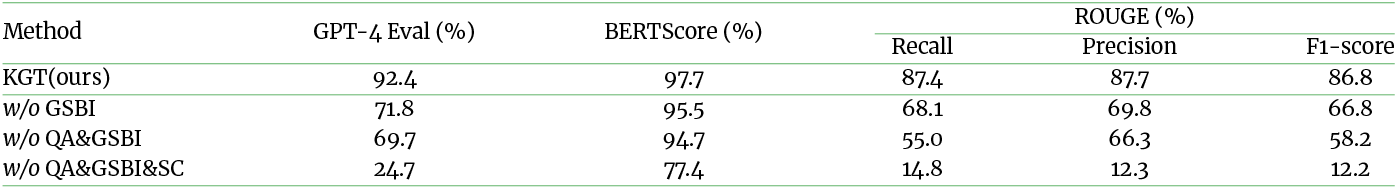
Ablation study of the KGT framework under Code-Llama-13B.

### Implementation Settings

Our knowledge graph is quite large, with a complex schema, and typically involves input tokens within 1300. Our experiment does not require fine-tuning, and the inference time is related to the model size and computational resources. For example, when using our method, KGT, with the Code-Llama-13B model on an 80GB A100 GPU, it occupies 33GB of VRAM. Without any acceleration frameworks, the inference requires four passes, each taking around 20 seconds.

### Case studies

#### Drug repositioning

Drug repositioning emerge as a promising strategy to accelerate the process of drug development. This approach involves identifying new therapeutic uses for existing drugs, thereby saving time and resources typically required for bringing a new drug to market [45]. Our system is capable of investigating the potential repositioning of carteolol for the treatment of hemangiomas. The example is shown in Supplementary Material Table.S2 and relational diagram is shown in Fig. 4(A). Utilizing the system’s knowledge graph, a relational chain is delineated, illustrating that propranolol, another inhibitor of ADRB1, is effectively employed in the treatment of hemangiomas. The system harnesses this insight to formulate a hypothesis that carteolol, by virtue of its similar mechanism of inhibition, could be potentially repositioning for treating hemangiomas [46]. This hypothesis would serve as a precursor to clinical trials and research, potentially expediting the availability of an additional therapeutic option for hemangiomas patients.

**Figure 4.**
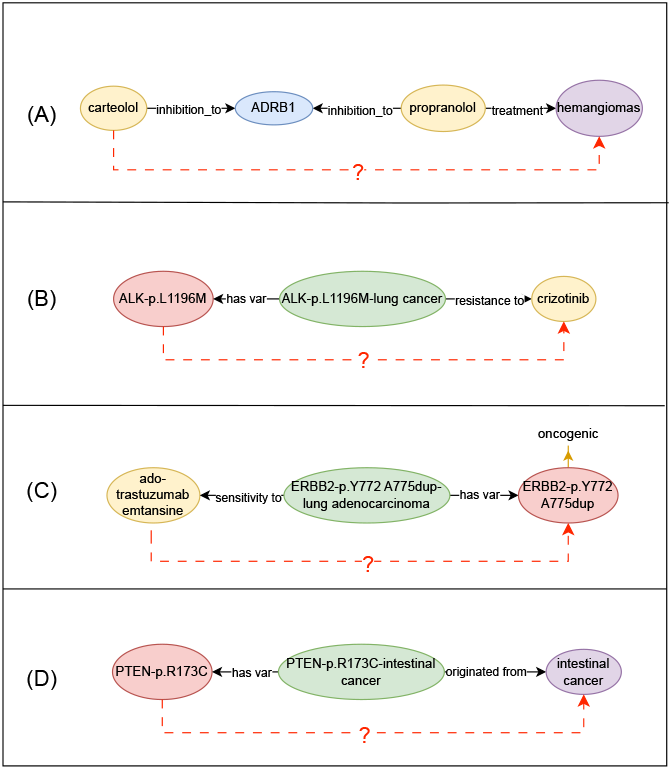
(A), (B), (C) and (D) respectively represent the relational diagrams of drug repositioning, drug resistance research, individualized treatmen and selection and understanding of biomarkers.

#### Drug resistance research

Drug resistance in cancer treatment poses a significant challenge in clinical oncology. Understanding the genetic basis of resistance can lead to more effective treatment strategies and personalized medicine approaches. Research in drug resistance involves determining why certain cancer carrying mutated gene are not responsive to specific drugs and finding ways to overcome this resistance [47]. Our system is capable of exploring drug resistance in cancer. The example is shown in Supplementary Material Table.S3 and relational diagram is shown in Fig. 4(B). The KG data indicates that the ALK-p.L1196M mutation, which is associated with gastric cancer, has a known resistance to Nalatinib [48, 49]. The LLM processes this information and infers that due to this resistance, Nalatinib might not be an effective medication for treating cancers caused by the ALK-p.L1196M mutation. The case highlights the critical importance of understanding specific gene-drug interactions in drug resistance research. It demonstrates how certain gene mutations could render a drug ineffective, which in turn could guide oncologists in choosing alternative treatments or developing new drugs that can bypass or target the resistance mechanisms. By accelerating the process of understanding drug resistance, these AI-driven systems can contribute to improved patient outcomes and the optimization of cancer treatment protocols.

#### Individualized treatment

Details on individualized treatment are provided in Supplementary Material Case studies A. It is important to note that this example is included solely to illustrate the technical capabilities of the proposed method. The output generated in this example has not been validated for clinical use, and further validation in clinical settings would be required before any such application.

#### Selection and understanding of biomarkers

Details on selection and understanding of biomarkers are provided in Supplementary Material Case studies B.

## Discussion

In this paper, we introduce a novel framework KGT, which employs LLMs for reasoning on the schema of KGs, to enhance the reasoning abilities of LLMs in areas with missing domain data by utilizing domain-specific knowledge graphs, such as oncology knowledge graphs, thereby addressing the issue of factual hallucinations in LLMs. Our method excels in extracting, validating, and refining factual knowledge throughout the LLMs’ reasoning process. It seamlessly integrates with various LLMs, including open-source models like Code-Llama, and enhances the capabilities of LLMs solely through prompt engineering and in-context learning, without any fine-tuning. This grants it significant generalizability.

We possess an extensive oncology knowledge graph and have established a benchmark based on it to evaluate the capabilities of various methods. When tested on PcQA using various open-source LLMs, the KGT framework performs exceptionally well, surpassing the current best methods by 33%. This significant improvement positions our approach as a pioneering benchmark in biomedical KGQA, setting a new standard that advances beyond previously established best practices. Additionally, through case studies, our approach has been shown to effectively provide therapeutic plans, generate valuable hypotheses for drug repositioning, identify potential drug targets, and study drug resistance. This underscores the practical value of the KGT framework in delivering insightful contributions that aid in the development and optimization of treatment strategies. Each case study’s conclusions are further validated by evidence from previously published research papers, enhancing the credibility and impact of our findings.

However, it is important to note that the constructed QA dataset and the corresponding published subset of the SmartQuerier Oncology Knowledge Graph (SOKG) were specifically designed to validate the effectiveness of the KGT framework within this study. While the dataset is highly relevant to biomedical applications, its scope is primarily focused on validating the proposed method. Therefore, it may not cover all potential use cases. Additionally, our system currently has the drawback of not performing fuzzy matching; if a drug name is misspelled by even one letter, it fails to retrieve information from the knowledge graph. Therefore, we plan to improve this aspect in the future to enhance the system’s usability and reliability. Our ultimate goal is to create a robust framework applicable to the rapidly evolving domain of medical knowledge, supporting healthcare professionals in delivering personalized, precise medication tailored to the individual needs of each patient.

Finally, we affirm that this study serves as a proof-of-concept, aiming to showcase the technical feasibility and initial efficacy of the method, which has not been validated in actual clinical practice. In any clinical or medical decision-making, reliance should always be placed on the judgment and guidance of professional healthcare practitioners.

## Availability of Source Code and Requirements

Project name: bioKGQA-KGT

- Project homepage: https://github.com/yichun10/bioKGQA-KGT.git.
- Operating system(s): Linux (Ubuntu)
- Resource usage in inference step: A Linux (Ubuntu) system with at least 2 CPU cores and 32 GB of VRAM. The GPU card needs at least 60GB VRAM(either two 32GB V100s or one 80GB A100).
- Programming language: Shell Script (Bash) with Python 3.10.13
- Other requirements: Python 3.10.13 with GPU/CPU support, neo4j 5.13.0 (please see more requirements on Github repository).
- Licenses: MIT license
- Research Resource Identifier (#RRID): SCR_025176

## Data Availability

We have publicly provided a subset of the SmartQuerier Oncology Knowledge Graph necessary for reproducing the research. An archival copy of the code and the subgraph of the Knowledge Graph used in this research is available via Software Heritage [33], and the code and datasets can be accessed via GitHub [50]. Additionally, the prompts used in interactions with LLMs [1, 2, 38, 42, 43, 44] during this research are available in the supplemental material. For access to the complete SmartQuerier Oncology Knowledge Graph data, please contact at.

## Supplementary material

Supplementary material is available at Supplementary material.pdf.

### Abbreviations

KG: knowledge graph
LLM: large language model
NLP: natural language processing
SFT: supervised fine-tuning
RLHF: reinforcement learning with human feedback
CF: catastrophic forgetting
CoT: Chain-of-thought
APE: automatic prompt engineer
KGQA: knowledge graph question answering
BFS: breadth-first search
PcQA: Pan-cancer Question Answering
ICL: in-context learning
GPT: generative pre-trained transformer.

## Ethical Statement

This study involves the generation of a biomedical question-answer dataset derived from a biomedical knowledge graph developed by our team. The knowledge graph has been meticulously constructed using non-personalized data obtained from various credible biomedical sources. The data collection and utilization processes strictly comply with all relevant legal regulations and ethical guidelines, ensuring the highest standards of data security and privacy. The dataset adheres rigorously to data protection principles and contains no sensitive personal information or identifiable individual health data. Furthermore, as the data collection and processing activities in this study do not involve human subjects, this research did not require ethical review or approval.

## Authors’ Contributions

Y.F. and L.Z. conceived the project. Y.F. proposed a KGQA benchmark, developed the KGT framework, implemented the code, conducted the experiments, and drafted the manuscript. C.M. contributed the SmartQuerier Oncology Knowledge Graph. Y.L. and L.Z. supervised the study. All authors read and approved the final manuscript.

## References

1. OpenAI, ChatGPT (Nov 30 version) [Large language model]; 2022. https://chat.openai.com/chat.

2. Touvron H, Martin L, Stone K, Albert P, Almahairi A, Babaei Y, et al. Llama 2: Open foundation and fine-tuned chat models [Large language model]. arXiv preprint 230709288 2023;.

3. Ji Z, Lee N, Frieske R, Yu T, Su D, Xu Y, et al. Survey of hallucination in natural language generation. ACM Computing Surveys 2023;55(12):1–38.

4. Liu T, Zheng X, Chang B, Sui Z. Towards faithfulness in open domain table-to-text generation from an entity-centric view. In: Proceedings of the AAAI Conference on Artificial Intelligence, vol. 35; 2021. p. 13415–13423.

5. Kang D, Hashimoto T. Improved natural language generation via loss truncation. arXiv preprint 200414589 2020;.

6. Pan S, Luo L, Wang Y, Chen C, Wang J, Wu X. Unifying large language models and knowledge graphs: A roadmap. IEEE Transactions on Knowledge and Data Engineering 2024;.

7. Han T, Adams LC, Papaioannou JM, Grundmann P, Oberhauser T, Löser A, et al. MedAlpaca–An Open-Source Collection of Medical Conversational AI Models and Training Data. arXiv preprint 230408247 2023;.

8. Yunxiang L, Zihan L, Kai Z, Ruilong D, You Z. Chatdoctor: A medical chat model fine-tuned on llama model using medical domain knowledge. arXiv preprint 230314070 2023;.

9. Singhal K, Azizi S, Tu T, Mahdavi SS, Wei J, Chung HW, et al. Large language models encode clinical knowledge. arXiv preprint 221213138 2022;.

10. Wang H, Liu C, Xi N, Qiang Z, Zhao S, Qin B, et al. Huatuo: Tuning llama model with chinese medical knowledge. arXiv preprint 230406975 2023;.

11. Xiong H, Wang S, Zhu Y, Zhao Z, Liu Y, Wang Q, et al. Doctorglm: Fine-tuning your chinese doctor is not a herculean task. arXiv preprint 230401097 2023;.

12. Zhang H, Chen J, Jiang F, Yu F, Chen Z, Li J, et al. HuatuoGPT, towards Taming Language Model to Be a Doctor. arXiv preprint 230515075 2023;.

13. Yang S, Zhao H, Zhu S, Zhou G, Xu H, Jia Y, et al. Zhongjing: Enhancing the chinese medical capabilities of large language model through expert feedback and real-world multi-turn dialogue. arXiv preprint 230803549 2023;.

14. Tian Y, Gan R, Song Y, Zhang J, Zhang Y. ChiMed-GPT: A Chinese Medical Large Language Model with Full Training Regime and Better Alignment to Human Preferences. arXiv preprint 231106025 2023;.

15. Ouyang L, Wu J, Jiang X, Almeida D, Wainwright C, Mishkin P, et al. Training language models to follow instructions with human feedback. Advances in Neural Information Processing Systems 2022;35:27730–27744.

16. Luo Y, Yang Z, Meng F, Li Y, Zhou J, Zhang Y. An empirical study of catastrophic forgetting in large language models during continual fine-tuning. arXiv preprint 230808747 2023;.

17. Li Z, Hoiem D. Learning without forgetting. IEEE transactions on pattern analysis and machine intelligence 2017;40(12):2935–2947.

18. Liu V, Chilton LB. Design guidelines for prompt engineering text-to-image generative models. In: Proceedings of the 2022 CHI Conference on Human Factors in Computing Systems; 2022. p. 1–23.

19. Wei J, Wang X, Schuurmans D, Bosma M, Xia F, Chi E, et al. Chain-of-thought prompting elicits reasoning in large language models. Advances in Neural Information Processing Systems 2022;35:24824–24837.

20. Zhou Y, Muresanu AI, Han Z, Paster K, Pitis S, Chan H, et al. Large language models are human-level prompt engineers. arXiv preprint 221101910 2022;.

21. Sun H, Dhingra B, Zaheer M, Mazaitis K, Salakhutdinov R, Cohen WW. Open domain question answering using early fusion of knowledge bases and text. arXiv preprint 180900782 2018;.

22. Sun H, Bedrax-Weiss T, Cohen WW. Pullnet: Open domain question answering with iterative retrieval on knowledge bases and text. arXiv preprint 190409537 2019;.

23. Zhang J, Zhang X, Yu J, Tang J, Tang J, Li C, et al. Subgraph retrieval enhanced model for multi-hop knowledge base question answering. arXiv preprint 220213296 2022;.

24. Chen Y, Wu L, Zaki MJ. Bidirectional attentive memory networks for question answering over knowledge bases. arXiv preprint 190302188 2019;.

25. Saxena A, Tripathi A, Talukdar P. Improving multi-hop question answering over knowledge graphs using knowledge base embeddings. In: Proceedings of the 58th annual meeting of the association for computational linguistics; 2020. p. 4498–4507.

26. Lan Y, He G, Jiang J, Jiang J, Zhao WX, Wen JR. A survey on complex knowledge base question answering: Methods, challenges and solutions. arXiv preprint 210511644 2021;.

27. Das R, Zaheer M, Thai D, Godbole A, Perez E, Lee JY, et al. Case-based reasoning for natural language queries over knowledge bases. arXiv preprint 210408762 2021;.

28. Jiang J, Zhou K, Zhao WX, Wen JR. Unikgqa: Unified retrieval and reasoning for solving multi-hop question answering over knowledge graph. arXiv preprint 221200959 2022;.

29. Jiang J, Zhou K, Dong Z, Ye K, Zhao WX, Wen JR. Structgpt: A general framework for large language model to reason over structured data. arXiv preprint 230509645 2023;.

30. Sun J, Xu C, Tang L, Wang S, Lin C, Gong Y, et al. Think-on-Graph: Deep and Responsible Reasoning of Large Language Model on Knowledge Graph. In: The Twelfth International Conference on Learning Representations;. .

31. LUO L, Li YF, Haf R, Pan S. Reasoning on Graphs: Faithful and Interpretable Large Language Model Reasoning. In: The Twelfth International Conference on Learning Representations;. .

32. Kim J, Kwon Y, Jo Y, Choi E. KG-GPT: A general framework for reasoning on knowledge graphs using large language models. arXiv preprint 231011220 2023;.

33. Feng Y, Zhou L, Ma C, Zheng Y, He R, Li Y, Knowledge Graph-based Thought: a knowledge graph enhanced LLMs framework for pan-cancer question answering (Version 1); 2024. [Computer software]. https://archive.softwareheritage.org/swh:1:dir:4d5d3acbd7784d97229a0a5ba0453f67f73ed6cf;origin=https://github.com/yichun10/bioKGQA-KGT;visit=swh:1:snp:1906dbbfc88c9d1c8b7acf7deb7495e8002cbafa;anchor=swh:1:rev:9a0244de046118fb6d2423912fd0b34df7fd052c.

34. Wang J, Wu M, Huang X, Wang L, Zhang S, Liu H, et al. SynLethDB 2.0: a web-based knowledge graph database on synthetic lethality for novel anticancer drug discovery. Database 2022;2022:baac030.

35. Zhu C, Yang Z, Xia X, Li N, Zhong F, Liu L. Multimodal reasoning based on knowledge graph embedding for specific diseases. Bioinformatics 2022;38(8):2235–2245.

36. Zhang Y, Dai H, Kozareva Z, Smola A, Song L. Variational reasoning for question answering with knowledge graph. In: Proceedings of the AAAI conference on artificial intelligence, vol. 32; 2018. .

37. Kim J, Park S, Kwon Y, Jo Y, Thorne J, Choi E. FactKG: Fact verification via reasoning on knowledge graphs. arXiv preprint 230506590 2023;.

38. Achiam J, Adler S, Agarwal S, Ahmad L, Akkaya I, Aleman FL, et al. GPT-4 Technical Report (Mar 14 version) [Large language model]. arXiv preprint 230308774 2023;.

39. Zhang T, Kishore V, Wu F, Weinberger KQ, Artzi Y. Bertscore: Evaluating text generation with bert. arXiv preprint 190409675 2019;.

40. Lin CY. Rouge: A package for automatic evaluation of summaries. In: Text summarization branches out; 2004. p. 74–81.

41. Dong Q, Li L, Dai D, Zheng C, Wu Z, Chang B, et al. A survey for in-context learning. arXiv preprint 230100234 2022;.

42. Roziere B, Gehring J, Gloeckle F, Sootla S, Gat I, Tan XE, et al. Code llama: Open foundation models for code [Large language model]. arXiv preprint 230812950 2023;.

43. Luo L, Ning J, Zhao Y, Wang Z, Ding Z, Chen P, et al. Taiyi: a bilingual fine-tuned large language model for diverse biomedical tasks [Large language model]. arXiv preprint 231111608 2023;.

44. Tunstall L, Beeching E, Lambert N, Rajani N, Rasul K, Belkada Y, et al., Zephyr: Direct Distillation of LM Alignment [Large language model]; 2023.

45. He S, Liu X, Ye X, Tetsuya S. Analysis of Drug Repositioning and Prediction Techniques: A Concise Review. Current Topics in Medicinal Chemistry 2022;22(23):1897–1906.

46. Gan Lq, Wang H, Ni Sl, Tan Ch. A prospective study of topical carteolol therapy in Chinese infants with superficial infantile hemangioma. Pediatric Dermatology 2018;35(1):121–125.

47. Gottesman MM. Mechanisms of cancer drug resistance. Annual review of medicine 2002;53(1):615–627.

48. Alshareef A, Zhang HF, Huang YH, Wu C, Zhang JD, Wang P, et al. The use of cellular thermal shift assay (CETSA) to study Crizotinib resistance in ALK-expressing human cancers. Scientific reports 2016;6(1):33710.

49. Simionato F, Frizziero M, Carbone C, Tortora G, Melisi D. Current strategies to overcome resistance to ALK-inhibitor agents. Current drug metabolism 2015;16(7):585–596.

50. Feng Y, Zhou L, Ma C, Zheng Y, He R, Li Y, bioKGQA-KGT: Knowledge Graph-based Thought; 2024. https://github.com/yichun10/bioKGQA-KGT.

